# Disease associated mutations in mitochondrial precursor tRNAs affect binding, m1R9 methylation and tRNA processing by mtRNase P

**DOI:** 10.1101/2020.07.07.191726

**Authors:** Agnes Karasik, Carol A. Fierke, Markos Koutmos

## Abstract

Mitochondrial diseases linked to mutations in mitochondrial(mt) tRNA sequences are abundant. However, the contributions of these tRNA mutations to the development of diseases is mostly unknown. Mutations may affect interactions with (mt)tRNA maturation enzymes or protein synthesis machinery leading to mitochondrial dysfunction. In human mitochondria, the first step of tRNA processing is the removal of the 5’end of precursor tRNAs (pre-tRNA) catalyzed by the three-component enzyme, mtRNase P. Additionally, one sub-component of mtRNase P, mitochondrial RNase P protein 1 (MRPP1) catalyzes methylation of the R9 base in pre-tRNAs. Despite the central role of 5’ end processing in mitochondrial tRNA maturation, the link between mtRNase P and diseases is mostly unexplored. Here we investigate how 11 different human disease-linked mutations in (mt)pre-tRNAs affect the activities of mtRNase P. We find that several mutations weaken the pre-tRNA binding affinity, while the majority of mutations decrease 5’ end processing and methylation activity catalyzed by mtRNase P. Furthermore, all of the investigated mutations in (mt)pre-tRNA^Leu(UUR)^ alter the tRNA fold which contributes to the partial loss of function of mtRNase P. Overall, these results reveal an etiological link between early steps of (mt)tRNA-substrate processing and mitochondrial disease.

## INTRODUCTION

Mitochondria serve as the powerhouse for the eukaryotic cell by synthesizing ATP. Due to its endosymbiotic origin, human mitochondria possess a circular genome (mtDNA) encoding 2 rRNAs, 22 tRNAs and 13 mRNAs that are essential for mitochondrial function. The mRNAs encode proteins involved in oxidative phosphorylation. Similar to the nuclear genome, mutation of mtDNA often leads to the development of diseases (Abbott et al., 2014; Chinnery, 2015; Finsterer and Kothari, 2014; Lightowlers et al., 2015; Schapira, 2012; Suzuki et al., 2011). One in every 4,000 children in the United States is born with a mitochondrial-related disease. Despite this, the pathogenesis of these diseases is not clearly understood and treatment is limited (Chinnery, 2015; Lightowlers et al., 2015; Schapira, 2012). Therefore, further research is needed to understand the underlying cause of these diseases and to develop new therapeutic strategies. Importantly, the majority (>50%) of mitochondrial disease-associated mtDNA mutations are located in mitochondrial transfer RNAs (tRNA) genes that occupy ~10% of the mtDNA (Abbott et al., 2014; Levinger et al., 2004a; Wittenhagen and Kelley, 2003). tRNAs are essential elements required for protein synthesis and they require several maturation steps before they become fully functional [9]. A link between mitochondrial diseases and impaired tRNA maturation in the mitochondria is beginning to emerge (Chatfield et al., 2015b; Levinger et al., 2004a; Levinger and Serjanov, 2012; Wang et al., 2011). The human mitochondrial genome encodes for three polycistronic units with tRNAs genes interspersed between rRNA and mRNA sequences (Ojala et al., 1981). These encoded polycistronic units contain no or only short non-coding sequences between tRNA genes and other coding regions, therefore processing of the 3’ and 5’ ends of tRNA processing also contributes to the maturation of rRNAs and mRNAs. As a consequence, improper tRNA processing can influence maturation of other essential RNAs in the mitochondria (Levinger et al., 2004a; Wittenhagen and Kelley, 2003). When RNA maturation is disturbed, the amount of mature RNAs available for normal function such as protein translation is decreased, potentially leading to mitochondrial dysfunction. Since mitochondria are the major source of energy of the cell, mitochondrial impairment contributes to the failure of physiological processes that require high amounts of ATP and therefore to development of disease.

In human mitochondria the first step of pre-tRNA maturation is the removal of the 5’ end of precursor tRNAs (pre-tRNA) (Sanchez et al., 2011). This early step is catalyzed by the three-component enzyme mitochondrial ribonuclease P (mtRNaseP) (Holzmann et al., 2008). In this complex, human MRPP3 (<u>M</u>itochondrial <u>R</u>Nase <u>P</u> <u>P</u>rotein 3 or <u>P</u>rotein <u>O</u>nly <u>R</u>Nase <u>P</u>, PRORP) harbors the catalytic core for 5’ end cleavage, however, it is only active in the presence of the sub-complex consisting of MRPP1 and MRPP2 (Mitochondrial RNase P Protein 1 and 2)(Holzmann et al., 2008). In addition to activating the 5’ end cleavage function of mtRNAse P, the MRPP1/2 sub-complex independently catalyzes methyl transfer to the ninth base (m1A9/m1G9) of human mitochondrial tRNAs (Vilardo et al., 2012). In this sub-complex, MRPP2 (17β-hydroxysteroid dehydrogenase type 1, SDR5C1) is proposed to serve as a scaffold protein supporting the S-adenosyl-methionine (SAM) dependent methyltransferase function of MRPP1 (TRM10C) (Vilardo et al., 2012).

mtRNase P is essential and plays a central role in mitochondrial function (Munch and Harper, 2016; Rackham et al., 2016; Sanchez et al., 2011; Sen et al., 2016). Disturbance of 5’ end pre-tRNA processing is indicated in several mitochondrial diseases, such us maternally inherited essential hypertension, mitochondrial myopathy, MELAS and HSD10 disease (Bindoff et al., 1993; Chatfield et al., 2015b; Deutschmann et al., 2014a; Falk et al., 2016; Li and Guan, 2010; Rossmanith and Karwan, 1998; Vilardo and Rossmanith, 2015; Wang et al., 2011). Mitochondrial diseases can be inherited in two ways, either by mendelian mutations in the nuclear genome or mutations in the mitochondrial genome. In particular, mutations in the nuclear encoded MRPP2 are linked to HSD10 disease (Amberger et al., 2016; Chatfield et al., 2015b; Deutschmann et al., 2014b; Vilardo and Rossmanith, 2015). Strikingly, most clinically-observed mutations in MRPP2 do not affect the dehydrogenase activity of MRPP2 rather they alter the methyltransferase and 5’ end cleavage activity of mtRNase P due to improper mtRNase P complex formation (Chatfield et al., 2015b; Deutschmann et al., 2014b; Falk et al., 2016). In addition, a single nucleotide polymorphism in the MRPP3 gene influences mitochondrial tRNA methylation patterns and potentially leads to differences in metabolism amongst individuals (Hodgkinson et al., 2014). Furthermore, disease linked mutations in human (mt)pre-tRNAs can also impair 5’ end pre-tRNA processing, although studies of such mutations are limited. In the most comprehensive study so far, it has been shown that the A4263G (mt)tRNA^Ile^ mutation linked to maternally inherited essential hypertension can affect 5’ end cleavage and methyltransferase activity of the human mtRNase P (Wang et al., 2011). This mutation decreases mature mitochondrial tRNA^Ile^ levels in patient cell lines suggesting that the perturbed processing of tRNA^Ile^ is likely an underlying cause of this disease.

Despite the potential link between mtRNase P and some other mitochondrial diseases there are very few studies addressing the direct role of mtRNase P in disease (Wang et al., 2011). Hence, here we have *in vitro* transcribed 11 human mitochondrial pre-tRNAs containing disease linked mutations and investigated their effects on binding, methylation and nuclease activity of human mtRNase P. We found that most of the investigated mitochondrial pre-tRNA mutations negatively affected the methylation and nuclease activity of the mtRNase P complex. Additionally, we have shown that a subset of these pre-tRNA mutations influenced proper tRNA folding and tertiary structure and this may account for the decrease in mtRNase P activity. Taken together, our study reveals that disease-linked mutations in (mt)tRNAs result in reduced mtRNase P activity *in vitro* and suggests that the contributions of these mutations to disease development may be related to modulated and disrupted mtRNase P function *in vivo*.

## RESULTS

### Disease-related mutations in human mitochondrial pre-tRNAs can influence substrate binding of mtRNase P

Pre-tRNA 5’ end cleavage is the first tRNA modification step in human mitochondria (Rackham et al., 2016; Reinhard et al., 2017; Sanchez et al., 2011), thus proper substrate recognition by mtRNase P is crucial for efficient RNA processing and editing. It has been long proposed that human mitochondrial tRNA mutations contribute to diseases by altering the structure of immature tRNAs and impairing recognition by processing and modification enzymes (Abbott et al., 2014; Kelley et al., 2000, 2001; Levinger et al., 2004a; Mustoe et al., 2015; Rossmanith and Karwan, 1998; Wittenhagen and Kelley, 2003; Yan et al., 2006; Yasukawa et al., 2000). However, how mutations affect binding to mtRNase P has not been evaluated thus far. In order to address this question we measured binding affinities (*K*_*D,app*_) of mtRNase P for pre-tRNAs containing a variety of disease–linked mutations through *in vitro* fluorescent binding assays.

We previously demonstrated that (mt)pre-tRNA^Ile^, (mt)pre-tRNA^Leu(UUR)^ and (mt)pre-tRNA^Met^ are substrates for mtRNase P when *in vitro* transcribed and unmodified (Karasik et al., 2019). Since these pre-tRNAs are also hotspots for mitochondrial disease mutations (Levinger et al., 2004b; Li and Guan, 2010; Li et al., 2009; Wang et al., 2013; Wang et al., 2011) we investigated *in vitro* binding affinities of their mutant versions to mtRNase P. A representative set of mitochondrial disease tRNA mutations located in functional regions of (mt)tRNAs were chosen for this study (Figure 1). The majority of the selected mutations are located in D- and T-loop and stem (tRNA elbow region; Figure 1D) while one maps to the acceptor stem and another one is found in the leader sequence and specifically in the first position in the 5’ end of the tRNA leader (N_−1_ position).

**Figure 1.**
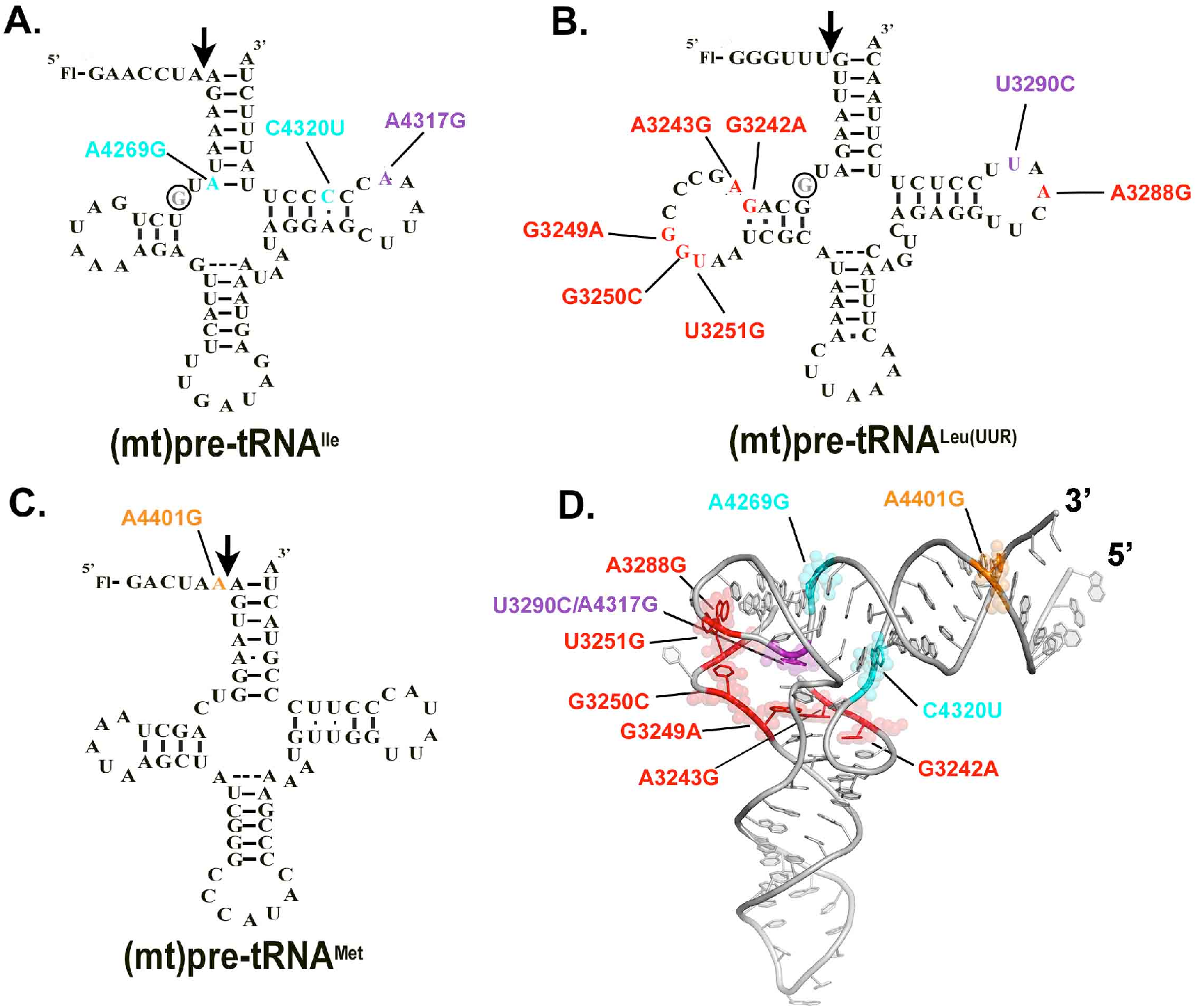
Locations of the investigated mitochondrial tRNA mutations in the predicted secondary structure of **A.** (mt)pre-tRNA^Ile^, **B.** (mt)pre-tRNA^Leu(UUR)^ and **C.** (mt)pre-tRNA^Met^. Circles indicate methylation sites for the MRPP1/2 sub-complex. **D.** Location of the mutations in the L-shaped tertiary structure of tRNAs. High-resolution structure of the investigated tRNAs are not available therefore we used a conserved eukaryotic tRNA^Phe^ with known structure (PDB ID: 1ehz) for generating the figure. Mutations are color coded according to the colors shown in Figure A., B. and C. Purple represents mutation at the same position in (mt)pre-tRNA^Leu(UUR)^ (U3290C) and (mt)pre-tRNA^Ile^ (A4317G).

We have *in vitro* transcribed and 5’ fluorescently labeled wild type and mutant (mt)pre-tRNAs and measured their binding affinity (*K*_*D,app*_) to mtRNase P based on previously established fluorescent polarization binding assays (Karasik et al., 2019; Liu et al., 2019). Only four of the (mt)pre-tRNA mutations moderately increase the *K*_*D,app*_ values by ~2-6 fold relative to the wild type (mt)pre-tRNA sequences (mutations A3243G G3249A, A4317G and C4320U) indicating reduced affinity to mtRNase P (Figure 2A, Supplemental Table 1.). However, several of the mutant (mt)pre-tRNAs (G3242A, G3250C, A3288G, U3290C, A4269G, A4401G) did not have any apparent effect on human mtRNase P binding affinity. This suggests that these disease-associated pre-tRNA mutations may influence cleavage or methyltransferase activity of mtRNase P or other mitochondrial functions, such as protein translation or activities of other mitochondrial tRNA modification enzymes, that potentially contributes to diseases.

**Figure 2.**
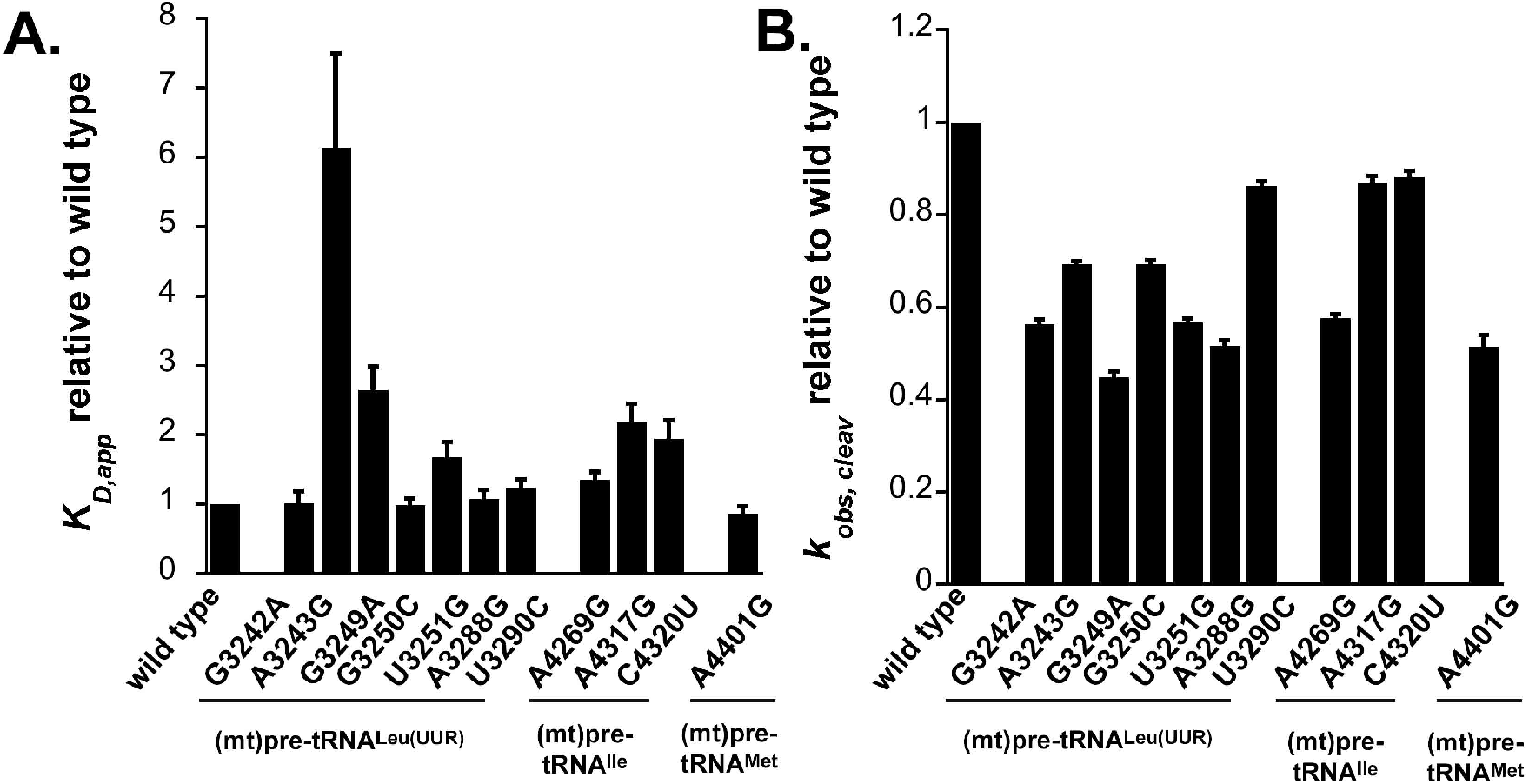
Disease-related tRNA mutations affect binding affinity and 5’ end pre-tRNA processing activity by the mtRNase P. **A.** Binding affinities of mutant mitochondrial pre-tRNAs for mtRNase P using standard binding conditions. 20 nM fluorescently labeled substrates were pre-incubated with 150 nM MRPP1/2 for 5 minutes and then titrated with 0-5 μM MRPP3 as described in (Karasik et al., 2019). Data were collected when binding reached equilibria from changes in fluorescenc anisotropy. The *K*_D_ values are calculated from a fit a binding isotherm to the data and reported values are divided by the wild type *K*_D_ values. **B.** Single turnover cleavage rates of mitochondrial pre-tRNAs containing mutations catalyzed by mtRNase P using standard cleavage conditions. Reactions contained 20 nM fluorescently labeled substrate, 1 μM MRPP3 and 0.4 μM MRPP1/2. The *k*_obs,cleavage_ values are calculated from a fit a single exponential to the data and reported values are divided by the wild type *k*_obs,cleavage_ values.

### Mitochondrial pre-tRNA mutations decrease 5’ end cleavage activity of mtRNase P

A mutation, A4263G, associated with inherited hypertension in (mt)pre-tRNA^Ile^ affected 5’ tRNA processing by the mtRNase P (Wang et al., 2011). Notably, even as little as ~30% decrease in mtRNase P activity can manifest in mitochondrial disease (Wang et al., 2011). Despite the connection between 5’ end tRNA processing and diseases in human mitochondria, studies of how mutant forms of mitochondrial tRNAs are processed by mtRNase P are limited. Thus, we investigated the effect of selected disease-associated mutations (Figure 1) in human mitochondrial pre-tRNAs on the 5’ end cleavage activity of mtRNase P. We employed a previously optimized fluorescent polarization based single turnover assay for mtRNase P that measures the reactivity of bound complex (mtRNase P•pre-tRNA) (Karasik et al., 2019). We found that the single turnover activity, as represented by the observed kinetic rate (*k*_*obs,cleavage*_), was reduced for all investigated tRNA mutants by 10% to 55% as compared to wild type (Figure 2B). The greatest impact on nuclease activity of mtRNase P was caused by (mt)pre-tRNA mutations, G3249A and A3288G, in D- and T- loops in (mt)pre-tRNA^Leu(UUR)^, respectively, and 4401G in the leader of (mt)pre-tRNAMet resulting in ≥50% decrease in activity. Additionally, we also observed a modest decrease (30-50%) in mtRNase P activity for mutations A3243A, G3242A, G3250C, U3251C in (mt)pre-tRNA^Leu(UUR)^ and A4269G in (mt)pre-tRNA^Ile^. However, some tRNA mutations, such as U3290C in (mt)pre-tRNA^Leu(UUR)^ and A3269A and A4317G in (mt)pre-tRNA^Ile^, only slightly affected (~10%) the 5’ pre-tRNA processing activity of mtRNase P. Taken together, several investigated mutations affected single turnover 5’ tRNA cleavage efficiency modestly and some severely of human mtRNase P.

### Disease-linked (mt)pre-tRNA^Leu(UUR)^ mutations alter m1G9 methylation activity in the mitochondria

The MRPP1/2 sub-complex is predicted to methylate the m1A9/m1G9 position of most mitochondrial tRNAs, however it is unknown if disease-linked mutations in pre-tRNAs can affect the activity of MRPP1/2. We have previously shown that both (mt)pre-tRNA^Leu(UUR)^ and (mt)pre-tRNA^Ile^ are methylated at their N9 position by mtRNase P based on single turnover primer extension methylation assays (Karasik et al., 2019). We therefore tested whether disease-linked mutations alter the methylation efficiency of MRPP1/2 for the various mutant versions of these two (mt)pre-tRNAs. Since the presence of human MRPP3 in methylation assays does not change the single turnover rate for (mt)pre-tRNA^Ile^ (Karasik et al., 2019; Vilardo et al., 2012), we assessed the effects of mitochondrial pre-tRNA mutations using the MRPP1/MRPP2 sub-complex without the presence of MRPP3 in the assays. A differential effect of mutations in (mt)pre-tRNA^Ile^ on MRPP1/2 methylation activity was observed. Specifically, MRPP1/2 single turnover methylation rates (*k*_*obs,meth*_) of both C4320U and A4317G (mt)pre-tRNA^Ile^ mutants displayed negligible differences as compared to the wild-type (mt)pre-tRNA^Ile^ (Figure 3A). In contrast, the presence of the A4269G mutation in (mt)pre-tRNA^Ile^ increased the MRPP1/2 single turnover methylation rate by ~2.5 fold (Figure 3A). Next, we investigated single turnover methylation rates for (mt)pre-tRNA^Leu(UUR)^ variants and mtRNase P. We previously found that the presence of MRPP3 in methylation assays enhances single turnover methylation rates of the MRPP1/2 sub-complex for the wild type (mt)pre-tRNA^Leu(UUR)^ and therefore we used equimolar concentrations of MRPP3 and the MRPP1/2 sub-complex in our methylation assays (Karasik et al., 2019). Most mutations in (mt)pre-tRNA^Leu(UUR)^ significantly decreased single turnover kinetics to such an extent that determining single turnover methylation rates were challenging. Therefore we measured the fraction of single turnover methylation at 1 hour compared to methylation of wild type (mt)pre-tRNA^Leu(UUR)^ (Figure 3B and C). All disease-linked (mt)pre-tRNA^Leu(UUR)^ mutants reduced methyltransferase activity significantly for mtRNase P (~40-95 %). However, we note that this may still be an underestimation of the effect of (mt)pre-tRNA^Leu(UUR)^ mutations on methyltranferase activity of mtRNase P since we compare their m1A9 methylation levels to fully methylated wild-type (mt)pre-tRNA^Leu(UUR)^. Taken together, all of the investigated disease linked tRNA mutations disrupt single turnover methyltransferase and/or cleavage activity of human mitochondrial mtRNase P.

**Figure 3.**
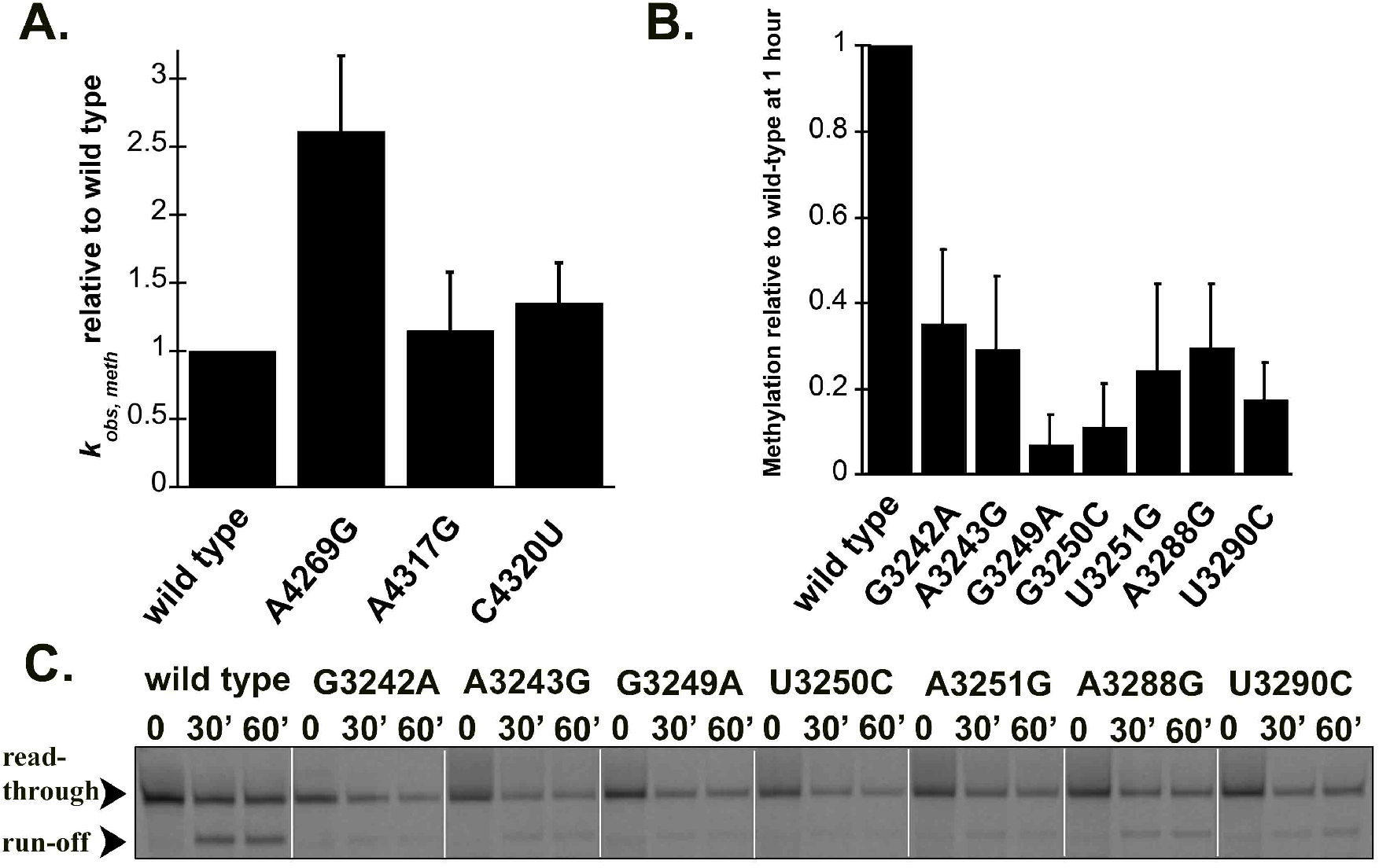
Disease-linked mutations in human (mt)pre-tRNA^Leu(UUR)^ decrease methylation efficiency catalyzed by the human mtRNase P complex. **A.** Single turnover methylation rates of mutant (mt)pre-tRNA^Ile^s relative to that of the wild type (mt)pre-tRNA^Ile^ catalyzed by the MRPP1/2 sub-complex using standard reaction conditions. Reaction contained 800 nM unlabeled pre-tRNA and 5 μM MRPP1/2. **B.** Quantification of methylation efficiency of mutant (mt)pre-tRNA^Leu(UUR)^s relative to wild type after 60 minutes incubation with mtRNase P. Reaction contained 800 nM unlabeled pre-tRNA and 5 μM MRPP1/2 and MRPP3. **C.** Representative gel showing methylation efficiencies of wild type and mutant (mt)pre-tRNA^Leu(UUR)^s by human mtRNase P at 30 and 60 minutes using standard reaction conditions. Black arrows indicate two products of primer extension reactions; the read through and the stop at the methylation sites.

### Mutations in mitochondrial pre-tRNAs can alter their folding and structural ensemble

Mutations in some mitochondrial tRNAs were shown to alter their structure which in turn may lead to decreased recognition by mitochondrial tRNA binding enzymes (such as the 3’ end cleavage enzyme RNase Z) (Levinger et al., 2004a; Levinger and Serjanov, 2012). Therefore, we also assessed the potential contribution of selected disease-related mutations in inducing changes to the structural ensemble and stability via UV melting experiments. In general, tRNAs exhibit several unfolding transitions upon denaturation, manifesting as hyperchromicity that can be monitored at 260 nM (Mustoe et al., 2015; Stein and Crothers, 1976). The first major transition is generally assigned to tertiary structure unfolding (Mustoe et al., 2015). Given that the tRNA structure, which is essential for efficient processing and functionality in protein synthesis, is stabilized by these tertiary contacts, we focused on dissecting this first transition of precursor (mt)tRNAs (Figure 4).

**Figure 4.**
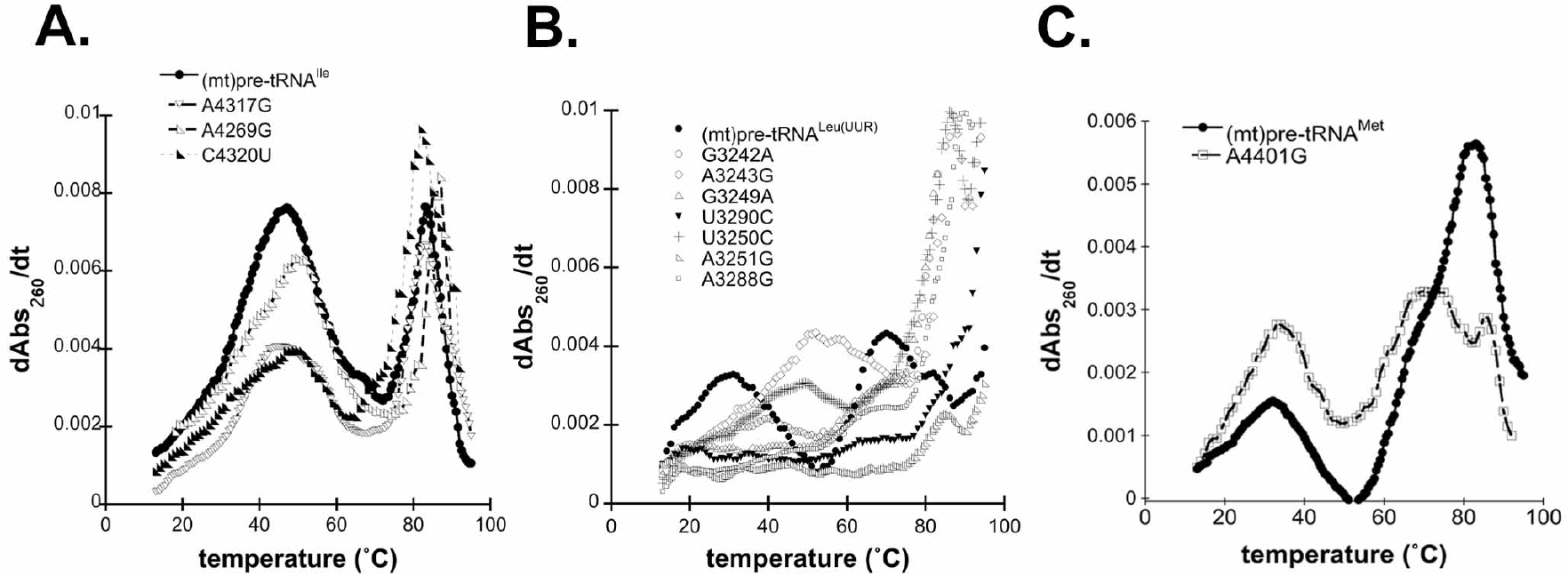
Disease-associated tRNA mutations alter the structure of pre-tRNAs. **A.** Representative UV melting profile curves for the wild type and mutant (mt)pre-tRNA^Ile^s. **B.** Representative UV melting profile curves of wild type and mutant (mt)pre-tRNA^Leu(UUR)^s. **C.** Representative UV melting profile curves of wild type and mutant (mt)pre-tRNA^Met^s.

Previous UV melting analysis of the wild type (mt)pre-tRNA^Ile^ revealed that two major transitions, with a melting temperature of 47.7 °C for the first transition, were observed (Karasik et al., 2019)(Figure 4A). Disease-associated tRNA mutations in (mt)pre-tRNA^Ile^ caused minor changes in the first observed transition as reflected by lower maximum derivates compared to that of the wild type (Figure 4A). In addition, while we observed one pronounced first transition for wild type (mt)pre-tRNA^Ile^ in the UV melting curve, the mutants appeared to have at least two separate transitions at lower temperatures (Supplemental Table 1). Although these transitions were overlapping and not well resolved, it is consistent with previous UV melting experiments carried out on wild type and disease-causing mutant (mt)tRNA^Ser^s, where introducing mutations into tRNA^Ser^ caused the separation of the pronounced first melting transition into two transitions with lower maximum derivates and unfolding enthalpy (Mustoe et al., 2015). These slight changes in the UV melting curves of (mt)pre-tRNA^Ile^ mutants demonstrate altered structural ensemble and folding pathways perhaps through disruption of interactions with Mg^2+^ or loss of H-bonding.

UV melting curves of wild type (mt)pre-tRNA^Leu(UUR)^ and (mt)pre-tRNA^Met^ exhibited at least three but at most four folding transitions (Figure 4B and C) (Karasik et al., 2019). However, we found that most (mt)pre-tRNA^Leu(UUR)^ mutants almost entirely lack the first transition observed for wild type (Figure 2B). This suggests that introducing these mutations into (mt)pre-tRNA^Leu(UUR)^ can severely affect folding and disrupt tertiary structure. For two of the (mt)pre-tRNA^Leu(UUR)^ mutants, A3243G and G3250C, we observed apparent first transitions at higher temperatures (~50°C). It is possible these particular mutations altered folding that allowed stronger interactions in the tertiary structure.

We have also examined wild type and A4401G (mt)pre-tRNA^Met^ in UV melting experiments. We found that the mutant (mt)pre-tRNA^Met^ exhibited similar first transition melting temperature as wild type (T_m A401G_=33.8 ± 1.1°C, T_m wild type_= 32.6 ± 1.1°C) albeit with higher maximum derivates and unfolding enthalpies (ΔH_A401G_= 27.2 ± 1.6, ΔH_wild type_= 24.0 ± 7.0 J) (Figure 4C). Mutation of this position also altered the second transition (assigned to unfolding of the secondary structure); we observe two distinct peaks at higher temperatures when A4401G is introduced. The observed differences between wild type and A4401G (mt)pre-tRNA^Met^ could be the consequence of base pairing between the N_−1_ and the 3’ discriminator base that influences overall folding and structure in mutant (mt)pre-tRNA^Met^.

## DISCUSSION

Reduced 5’ end processing by mtRNase P in the human mitochondria is linked to disease-related mutations in components of mtRNase P and mitochondrial tRNA genes (Chatfield et al., 2015a; Deutschmann et al., 2014b; Falk et al., 2016; Vilardo and Rossmanith, 2015; Wang et al., 2011). While mitochondrial tRNA gene mutations leading to diseases are numerous, there is only one extensive study investigating the connection between those and mtRNase P (Wang et al., 2011). Thus, we speculated that many more tRNA mutations in mtDNA linked to different diseases could affect mtRNase P function in the mitochondria. Here, we show that eleven mutations associated with distinct mitochondrial diseases in (mt)pre-tRNA genes significantly reduce binding, methylation and 5’ end processing activity of mtRNase P and influence pre-tRNA folding and structure that may account for some of the decreased enzymatic activities of mtRNase P *in vitro*.

Selected mutations in (mt)pre-tRNA^Leu(UUR)^ are linked to several diseases with a wide range of disease outcomes (Table 1), (Gattermann et al., 2004; Gerber et al., 2010; Goto et al., 1990; Goto et al., 1992; Li and Guan, 2010; Seneca et al., 2001; Sweeney et al., 1993; Wang et al., 2013; Wortmann et al., 2012). Seven of the eleven selected mitochondrial tRNA mutations are located near or in the elbow region of human mitochondrial (mt)pre-tRNA^Leu(UUR)^ (Figure 1) that was shown to be an important region for substrate recognition in plant PRORP homologs (Gobert et al., 2013; Imai et al., 2014; Klemm et al., 2017; Pinker et al., 2017). Therefore, these mutations have the potential to directly impair specific tRNA-mtRNase P interactions or influence the structural integrity of the elbow region of the pre-tRNA affecting proper tRNA binding position or orientation by mtRNase P. For instance, A3243G and G3249A mutations could potentially disrupt predicted stacking and H-bonding interactions, respectively, between the T- and D-loop (Suzuki et al., 2011). In our biochemical assays, we found that all investigated (mt)pre-tRNA^Leu(UUR)^ mutations severely impacted single turnover rates for m1G9 methylation (~40-95% as compared to wild type) and decreased single turnover 5’ end cleavage rates of mtRNase P by ~10-55%. However, these mutations in general had a smaller effect on the binding affinity, indicating that the structural changes affected the reactivity of the bound mtRNase P•mtpre-tRNA complex. This suggests that structural changes in mt-pre-tRNA caused by mutations are retained in the enzyme-bound complex. Consistent with this, all mutant (mt)pre-tRNA^Leu(UUR)^s show significantly altered UV melting profile as compared to the wild type referring to the affected ensemble of these mutant pre-tRNAs. Previously, we found that the wild type (mt)pre-tRNA^Leu(UUR)^ melting temperature (T_m_=~28°C) is significantly lower than human mitochondrial physiological temperature suggesting that this pre-tRNA is structurally unstable in the absence of other stabilizing factors, such as other protein partners or RNA modifications (Karasik et al., 2019). Our data suggest that the investigated mutations cause disturbances in folding and/or tertiary structure of pre-(mt)tRNA^Leu(UUR)^, as indicated by changes in the first transition observed in UV melting experiments. The severely affected folding and structure of investigated (mt)pre-tRNA^Leu(UUR)^s could explain the observed reduced 5’ end cleavage and methylation single turnover rates for mtRNase P. However, we found that binding of these mutant (mt)pre-tRNA^Leu(UUR)^s to mtRNase P (titrated for PRORP) was not altered significantly, except for A3243G (mt)pre-tRNA^Leu(UUR)^. This suggests that the altered structural properties of the tRNA^Leu(UUR)^ variants only minimally affect substrate recognition. We propose that although most of these mutants still bind to mtRNase P, they are not oriented properly in the active sites of MRPP1 and PRORP for optimal methyltransferase and cleavage activity. Overall, our data suggest that these disease-related mutations in (mt)pre-tRNA^Leu(UUR)^ can affect several functions of mtRNase P *in vitro* and potentially influence mitochondrial tRNA maturation pathway *in vivo*, however this needs further testing (Figure 5).

**Table 1.** Disease-linked tRNA mutations affect binding and cleavage rates of mtRNase P.

| human (mt)pre-tRNA | description of disease | references | $K_{D,app}$ (nM) | $k_{obs,cleavage}$ (min<sup>-1</sup>) | $k_{obs,meth}$ (min<sup>-1</sup>) |
| --- | --- | --- | --- | --- | --- |
| (mt)pre-tRNA<sup>Leu</sup><sub>(UUR)</sub> |  |  | 50.6 ± 5.3 | 0.309 ± 0.002 | 0.201 ± 0.040 |
| G3242A | cardiomyopathy and ineffective hematopoiesis | (Gattermann et al., 2004; Munch and Harper, 2016; Wortmann et al., 2012) | 51.3 ± 8.3 | 0.174 ± 0.002 | 0.115 ± 0.040 |
| A3243G | MELAS (mitochondrial encephalomyopathy, lactic-acidose and stroke-like episodes) and diabetes with deafness | (Gerber et al., 2010; Goto et al., 1990) | 310.4 ± 69.0 | 0.214 ± 0.003 | <0.1* |
| G3249A | Kearns-Sayre syndrome | (Seneca et al., 2001) | 133.3 ± 17.5 | 0.138 ± 0.004 | <0.1* |
| G3250C | mitochondrial myopathies | (Goto et al., 1992) | 49.6 ± 4.9 | 0.214 ± 0.003 | <0.1* |
| U3251G | mitochondrial myopathies | (Sweeney et al., 1993) | 84.2 ± 11.5 | 0.175 ± 0.003 | <0.1* |
| A3288G | mitochondrial myopathies | (Wang et al., 2013) | 53.9 ± 6.9 | 0.159 ± 0.004 | <0.1* |
| U3290C | possible hypertension factor | (Zhu et al., 2009a) | 62.0 ± 6.5 | 0.266 ± 0.003 | <0.1* |
| (mt)pre-tRNA<sup>Ile</sup> |  |  | 14.6 ± 1.6 | 0.573 ± 0.008 | 0.070 ± 0.015 |
| A4269G | fatal infantile cardiomyopathy (FICP) | (Taniike et al., 1992; Yasukawa et al., 2000) | 19.4 ± 1.8 | 0.326 ± 0.005 | 0.182 ± 0.023 |
| A4317G | fatal infantile cardiomyopathy (FICP) | (Zhu et al., 2009a) | 31.6 ± 3.9 | 0.493 ± 0.008 | 0.080 ± 0.013 |
| C4320U | early-onset severe encephalomyopathy | (Santorelli et al., 1995) | 28.1 ± 3.9 | 0.499 ± 0.008 | 0.094 ± 0.022 |
| (mt)pre-tRNA<sup>Met</sup> |  |  | 707 ± 168 | 0.286 ± 0.006 | N.A. |
| A4401G | hypertension | (Zhu et al., 2009b) | 606.7 ± 72.8 | 0.147 ± 0.007 | N.A. |

\* Due to the low methylation activities for these (mt)pre-tRNA<sup>Leu(UUR)</sup>s exact methylation rates cannot be derived.
N.A.=(mt)pre-tRNA does not harbor a methylation site

**Figure 5.**
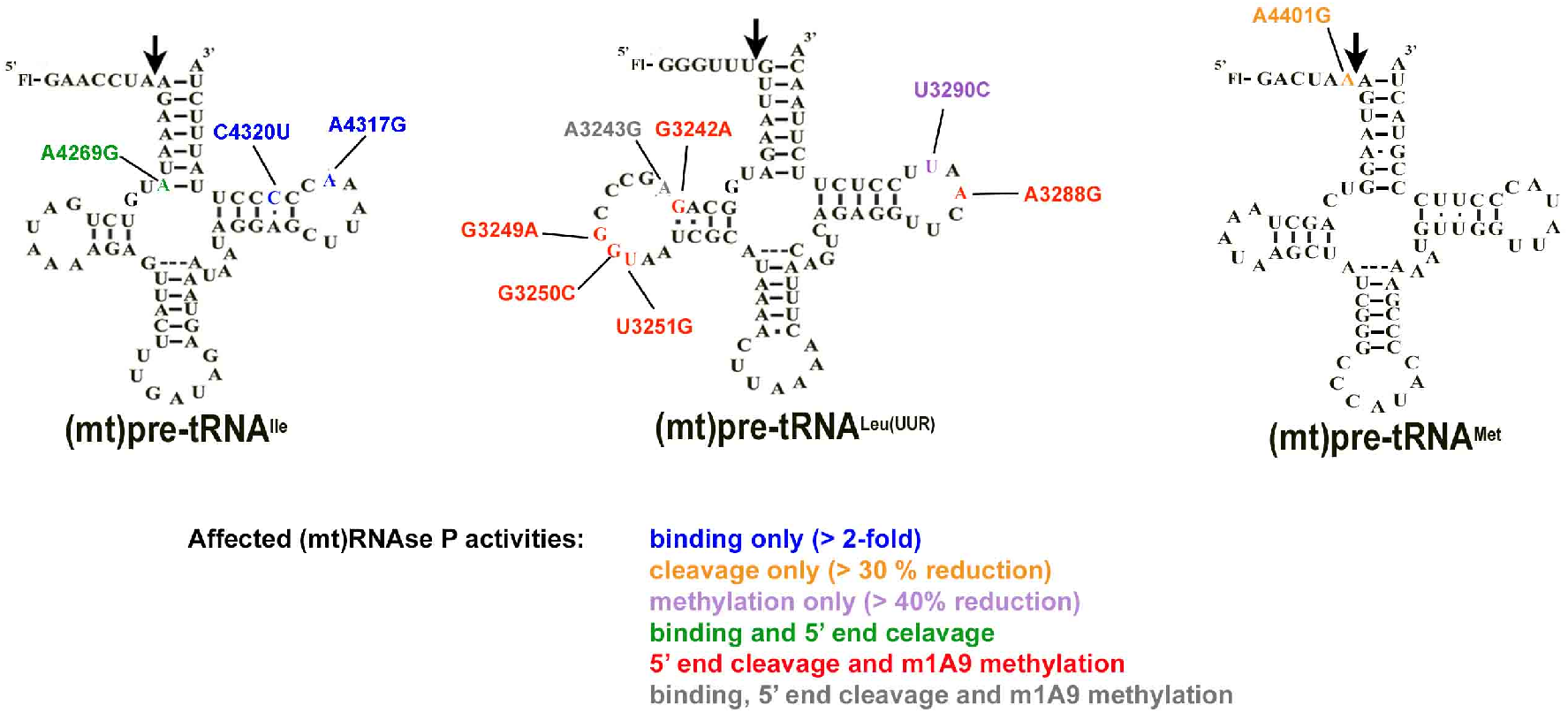
Summary of affected mtRNase P activities of each (mt)pre-tRNA mutations in this study.

Mitochondrial tRNA^Ile^ gene is also a hotspot for mutations; A4269G and A4317G mutations are associated with fatal infantile cardiomyopathy (FICP) (Hino et al., 2004; Taniike et al., 1992; Yasukawa et al., 2000), and C4320U (mt)pre-tRNA mutation is associated with early-onset severe encephalomiopathy (Table 1) (Santorelli et al., 1995). Thus, we also investigated these three (mt)pre-tRNA^Ile^s containing disease-associated mutations located near or in the elbow region of the pre-tRNA. It has been previously shown that the A4269G mutation in mature (mt)tRNA^Ile^, associated with the mitochondrial disease FICP, reduces (mt)tRNA^Ile^ aminoacetylation, *in vivo* half-life and *in vitro* melting temperature (Yasukawa et al., 2000). In agreement with these findings, we have observed that this mutation affected the folding and structure of (mt)pre-tRNA^Ile^ based on UV melting experiments and significantly reduced 5’ end pre-tRNA cleavage, however, it did not influence substrate affinity by mtRNase P based on our binding assays. The A4269G mutation is expected to disrupt local secondary tRNA structure by eliminating base pairing at the base of the acceptor stem and consequently cause changes in the structure of this region. Therefore, integrity in this area may play a role in proper orientation of the substrate for 5’ end cleavage in the RNase P•substrate complex. On the other hand, single turnover methylation rates for this mutant pre-tRNA were enhanced (~2.5 fold). This can be potentially attributed to better access to the methylation site by MRPP1 due to a more open structural arrangement of the acceptor stem. We also found that A4317G and C4320U mutant (mt)pre-tRNA^Ile^s were bound less tightly to mtRNase P and had slightly altered UV melting profiles compared to the wild type pre-tRNA^Ile^. However, these mutations did not influence single turnover methylation rates, and decreased 5’end cleavage activity of mtRNAs P by only ~10%. This suggests that the structural changes in these mutations are not recapitulated in the enzyme-bound complex. Furthermore, these data indicate that these mutations may affect different functions in the mitochondria than 5’ end processing or R9 methylation of pre-tRNAs. Nevertheless, we note that previous studies found that a ~ 20% decrease in single turnover activity of mtRNase P *in vitro* for A4263G (mt)pre-tRNA^Ile^ mutant led to significant accumulation of 5’ uncleaved mitochondrial pre-tRNAs *in vivo* (Wang et al., 2011). Therefore, the possible involvement of mtRNAse P in diseases linked to A4317G and C4320U tRNA mutations cannot be ruled out.

Plant homologs of MRPP3 have been show to interact with the N_−1_ base in the 5’ end leader of pre-tRNAs (Howard et al., 2016; Klemm et al., 2017). Therefore, we hypothesized that disease-linked mutations in this position could influence 5’end tRNA cleavage in the mitochondria that could contribute to the development of diseases. Hence we also investigated the effects of mutation in mitochondrial (mt)pre-tRNA^Met^ associated with hypertension (A4401G, found at the N_−1_ position) on mtRNase P activities (Table 1) (Zhu et al., 2009b). We found that this mutation affected 5’end cleavage, but not binding of mtRNase P. We posit that this mutation may influence the interaction between the 5’ end leader and MRPP3 by disrupting the proper orientation of the 5’ end leader bound to the active site of MRPP3, subsequently causing decreased 5’ end processing rates by mtRNase P. Since the folding and structure of this mutant pre-tRNA is also affected based on our UV melting data, subsequent steps in the mitochondrial tRNA maturation pathway are expected to also be influenced. Therefore, reduced 5’ end pre-tRNA processing could be one of the factors leading to hypertension in patients carrying A4401G mutation, however this needs further testing.

Freshly transcribed mitochondrial polycistronic units punctuated by tRNAs are expected to be subject to a hierarchical cascade of RNA processing and modifications. The 5’ end pre-tRNA processing and methylation by mtRNase P are first in line after transcription and required for downstream processes (Reinhard et al., 2017; Sanchez et al., 2011). Therefore, many pre-tRNA mutations could affect and decrease mtRNase P function first before they can also impact other downstream processes. Additional negative effects of these mutations in downstream processes and enzymes are possible and the impact on the latter may compound the initial effect on mtRNase P activity multiplying the adverse consequences of these mutations. In support, there is accumulating evidence that downstream enzymes from mtRNase P in the mitochondrial tRNA maturation pathway also have reduced activities with pre-tRNAs containing disease-linked mutations (Levinger et al., 2003; Levinger et al., 2004a; Levinger et al., 2004b; Levinger and Serjanov, 2012; Sissler et al., 2004; Yan et al., 2006). In particular, some of the investigated mutations (such as A4317G in (mt)pre-tRNA^Ile^ (Levinger et al., 2003)) have been previously shown to have inhibitory effect on 3’ tRNA maturation by human mitochondrial ELAC2 (Levinger et al., 2003; Levinger et al., 2004b). We also observed differential effects on enzyme activities (5’ end processing and methylation) for the same pre-tRNA mutation. Therefore, it is likely that tRNA mutations in the mtDNA affect several steps of pre-tRNA maturation and development of diseases could be the consequence of these additive sequential effects on enzyme activity. There is a wide range of manifestation of mitochondrial diseases associated with tRNA mutations that may arise from differences in how severely these mutations affect the different steps in the tRNA maturation pathway. However, *in vivo* studies are needed to investigate the exact role of mtRNase P in human mitochondrial diseases and the potential of the additive effects on the mitochondrial tRNA maturation pathway.

Taken together, we have found that the *in vitro* activities of mtRNase P are impacted when mtRNase P encounters pre-tRNAs carrying human disease-linked mutations (Figure 5). The observed impairment in mtRNase P activities could arise from changes in pre-tRNA structure when mutations are introduced. Our work potentially implicates the mtRNase P complex as a significant factor in the development of several mitochondrial diseases. Further understanding of the exact role of mtRNase P in the context of mitochondrial diseases is needed that may enable of development of new therapeutics in the future.

## MATERIALS AND METHODS

### *In vitro* transcription and 5’ end labeling of pre-tRNAs

5’ end fluorescein labeled wild-type and mutant pre-tRNAs were prepared as previously described (Karasik et al., 2019) by *in vitro* transcription (Howard et al., 2012; Karasik et al., 2016). Pre-tRNA DNAs were commercially synthesized by Integrated DNA Technologies and used as a template for *in vitro* transcription reaction. The reaction contained 50 mM Tris-HCl pH 8.0, 4 mM MgCl_2_, 1 mM spermidine, 5 mM dithiothreitol (DTT), 4 mM ATP, 4 mM CTP, 4 mM UTP, 1mM GTP, 4 mM guanosine- 5’-O-monophosphorothioate (GMPS), 350 μg/ml purified T7 RNA polymerase, 0.4 nmole DNA template containing T7 promoter. After 4 hours incubation at 37 °C, the reaction was stopped, the pre-tRNAs were 5’ end labeled and subsequently gel-purified as described before (Howard et al., 2012; Karasik et al., 2016). Unlabeled pre-tRNAs were made similarly to labeled ones, except transcription reaction contained 4 mM of GTP and excluded GMPS in addition to omitting the 5’ end labeling step with fluorescein. The concentrations of total and labeled pre-tRNA were measured with absorbance using a Nanodrop (Thermo Scientific) spectrophotometer and calculated by using the following extinction coefficients: 909,100 cm^−1^ M^−1^ for (mt)pre-tRNA^Ile7:0^, 927,300 cm^−1^ M^−1^ for (mt)pre-tRNA^Leu(UUR)6:0^, 838,200 cm^−1^ M^−1^ for (mt)pre-tRNA^Met6:0^.

### Protein expression and purification

Full length MRPP2 and His_6_-Δ39 MRPP1 lacking the predicted mitochondrial targeting sequences were cloned into pCDFDuet-1 (Novagen) vector (Liu et al., 2019). MRPP1 and 2 were co-expressed and purified together from *E. coli* (BL21 or Rosetta). First, expression was induced by addition of 200 μM isopropyl β-D-1-thiogalactopyranoside (IPTG) to transfected bacteria, after incubation overnight at 18°C. Bacterial cultures were collected and lysed as described before (Karasik et al., 2019; Karasik et al., 2016; Liu et al., 2019). The MRPP1/MRPP2 complex was purified using a nickel affinity column (GE Healthcare) and the His-tag was removed from MRPP1 by by incubation with TEV protease. MRPP1 and 2 formed a complex of ~150 kDa that was further purified by gel-filtration (Superdex 200, GE Healthcare). His_6-_Δ95 MRPP3 cDNA lacking the N-terminal disordered region was cloned into pMCSG7 vector and expressed and purified, as previously described (Karasik et al., 2019). In brief, MRPP3 expression was induced by addition of 200 μM IPTG to transfected *E. coli* BL21. Cells were harvested and lysed after overnight incubation similarly to the MRPP1/2 sub-complex. MRPP3 was purified by nickel affinity column and His-tag was removed by TEV-protease. Purity of the co-expressed MRPP1/2 sub-complex and MRPP3 was evaluated by SDS-PAGE (Sodium-dodecyl-sulfate polyacrylamide gel electrophoresis) and coomassie staining. Protein concentrations were measured by Nanodrop spectrophotometer (Thermo scientific) using 56.4 kDa as molecular weight and ε_280_ = 82,500cm^−1^ M^−1^ for MRPP3. The ratio of MRPP1/2 subunits as prepared is 1:4 based on analytical ultracentrifugation experiments (Liu et al., 2019). Therefore, the MRPP1/2 protein concentration was measured using 149 kDa as the molecular weight and extinction coefficient of 79,520 cm^−1^ mol^−1^.

### 5’ end cleavage single turnover cleavage assays

Pre-tRNA substrates were re-folded before each 5’ pre-tRNA cleavage reaction. First, pre-tRNAs were heated up to 95 °C for 3 minutes, cooled down to room temperature (~10-15 minutes), then folded by incubation in “cleavage reaction buffer” (30 mM MOPS pH 7.8, 150 mM NaCl, 1 mM MgCl_2_, 1 mM DTT) containing 1 mM Mg^2+^ for an additional 5 minutes at room temperature. All single turnover reactions were conducted as previously reported at 28 °C and saturating enzyme concentrations (Karasik et al., 2019). The assay temperature was chosen to be equal or be above the previously observed UV melting transitions (carried out under similar circumstances) to ensure proper folding of pre-tRNAs (Karasik et al., 2019). To initiate the reaction, a mixture of 500 nM MRPP1/MRPP2 complex and 100 nM to 2 μM Δ95 MRPP3 in cleavage reaction buffer was added to 20 nM fluorescently labeled pre-tRNA substrate. Changes in fluorescent polarization was followed at 28 °C using the ClarioStar plate reader (BMG LabTech). Data from at least three independent experiments were analyzed using Kaleidagraph 4.1.3. Eq 1. was fit to the data to calculate the observed rate constant (*k*_obs, cleavage_) and the standard error.

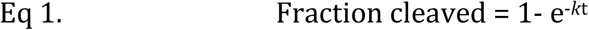

### Fluorescent polarization binding assays

Pre-tRNAs used in these experiments were folded as described above, except buffers used for folding contained 1 mM Ca^2+^ instead of Mg^2+^. All binding assays were performed in 30 mM MOPS pH 7.8, 150 mM NaCl, 6 mM CaCl_2_, 1 mM DTT (“binding buffer”) at 28 °C (Karasik et al., 2019). First, we pre-incubated 20 nM fluorescently labeled and folded pre-tRNAs with recombinant 150 nM MRPP1/2 to reach equilibria. Then we added increasing concentrations of MRPP3 (0-5 μM) and measured the fluorescent anisotropy values after an additional 5 min incubation. Data were corrected with the anisotropy measured in the absence of the protein but presence of 20 nM Fl-pre-tRNA. The binding isotherm (Eq. 2) was fit to the fluorescence polarization data from at least three independent experiments. Kaleidagraph 4.1.3 software was used to analyze data to calculate the dissociation constant (*K*_*D*_) and the standard error. In Eq 2 A is the observed anisotropy, A_o_ is the initial anisotropy, ΔA is the total change in anisotropy, P is the concentration of MRPP3, and *K*_D_ is the apparent dissociation constant.

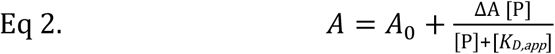

### Primer extension methylation assays

Unlabeled pre-tRNAs were prepared by *in vitro* transcription and gel-based purification. Before each experiment pre-tRNAs were folded in 1 mM Ca^2+^ containing “binding buffer”. Primer extension methylation assays were carried out as described before (Karasik et al., 2019). In short, 800 nM folded pre-tRNAs were incubated together with near-saturating (5 μM) MRPP1/2 or the reconstituted mtRNase P complex in “binding buffer” and 25 μM S-adenosyl methionine as described before (Karasik et al., 2019). Reaction was stopped at different time points by heating the sample for 30 seconds at 95 °C then placing it on ice. Terminated methyltransfer reactions were subjected to primer extension followed by separation of products by urea-gel electrophoresis (Karasik et al., 2019). Sequences for primers that bind to the anticodon loop were the following for wild type and mutant tRNA^Ile^, tRNA^Leu(UUR)^ and tRNA^Met^ respectively: 6-FAM-CTATTATTTACTCTATC; 6-FAM-CCTCTGACTGTAAAG; 6-FAM-CGGGGTATGGGCCCG. Data from at least three independent experiments were analyzed using Kaleidagraph 4.1.3. Eq 1 was fit to the data to calculate the single turnover rate constants (*k*_*obs, meth*_) and the standard error.

### UV melting experiments

Pre-tRNAs were folded as described above before each experiment using a “UV melting buffer” of final concentration of 10 mM PIPES pH 7, 150 mM NaCl, 1 mM MgCl_2_ and 1 μM pre-tRNA. Thermodynamic curves were obtained by heating the samples at 1 °C/min from 15 °C to 95 °C while absorbance at 260 nm was monitored at every 0.5 or 1 °C using a Cary 300 spectrophotometer. Cooling from 95 °C to 25 °C (hysteresis) resulted in a comparable curve to that obtained by heating. Data were corrected with data from blank measurements and analyzed with fitUVData.py and Global Melt Fit (Draper, 2000; Mustoe et al., 2015). Most pre-tRNA showed three or four transitions in agreement with previous findings (Mustoe et al., 2015). Average T_m_s and standard errors were calculated from at least three independent experiments and bootstrapping analysis was performed to ensure the robustness of the data where it was applicable.

## ACKNOWLEDGEMENTS

We thank Dr. Aranganathan Shamuganathan for supplying some of the MRPP1/2 enzyme preparation. We also thank Dr. Adrian Ferre-D’Amare, Dr. Robert Trachman and Dr. Xin Liu for providing great help for UV melting experiments. In addition we are greatful to Dr. Robert Trachman for reading the manuscript and providing valuable feedback. Special thanks to Meredith Purchal for her help in performing densitormetry.

