## Supplemental for "Disease associated mutations in mitochondrial precursor tRNAs affect binding, m1R9 methylation and tRNA processing by mtRNase P"

**Supplemental Table 1.** Melting temperature values for first transitions for wild type (mt)pre-tRNAs and (mt)pre-tRNA$^{Ile}$ mutants.

| human mitochondrial pre-tRNA | $T_m$1 (˚C) | $T_m$2 (˚C) |
|---|---|---|
| (mt)pre-tRNA$^{Leu(UUR)}$ | 28.7 ± 0.1 | - |
| (mt)pre-tRNA$^{Met}$ | 32.6 ± 1.1 | |
| (mt)pre-tRNA$^{Ile}$ | 47.7 ± 1.6 | |
| A4269G | 40.4 ± 5.4 | 49.3 ± 1.0 |
| A4317G | 42.5 ± 0.9 | 56.2 ± 0.5 |
| C4320T | 27.7 ± 3.7 | 47.5 ± 0.4 |

Supplemental Figure 1.

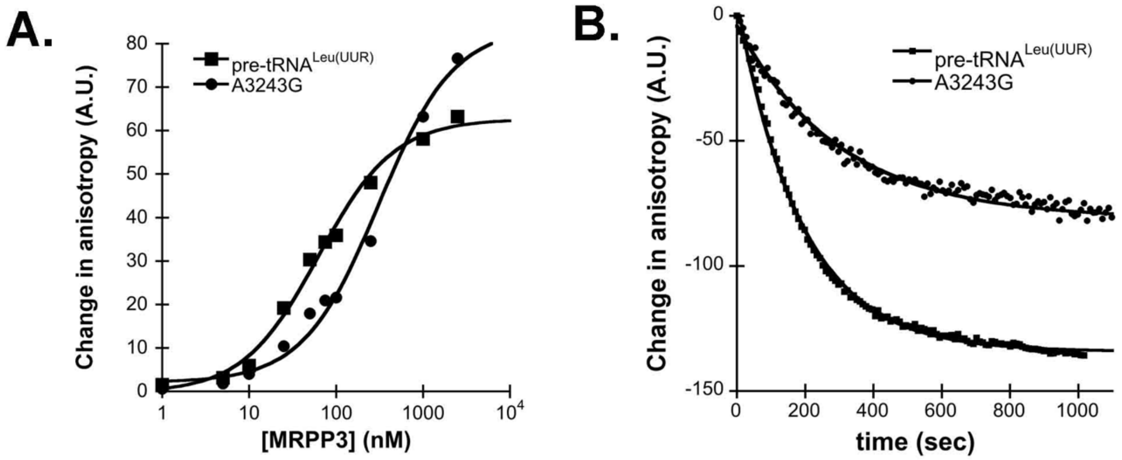


Supplemental Figure 1. A. Example of a wild type and a mutant (mt)pre-tRNA binding in the function of MRPP3 concentrations when 150 nM MRPP1/2 is present using standard binding assay conditions. B. Example of the single turnover cleavage of a wild type and a mutant (mt)pre-tRNA using standard cleavage assay conditions.

Supplemental Figure 2.

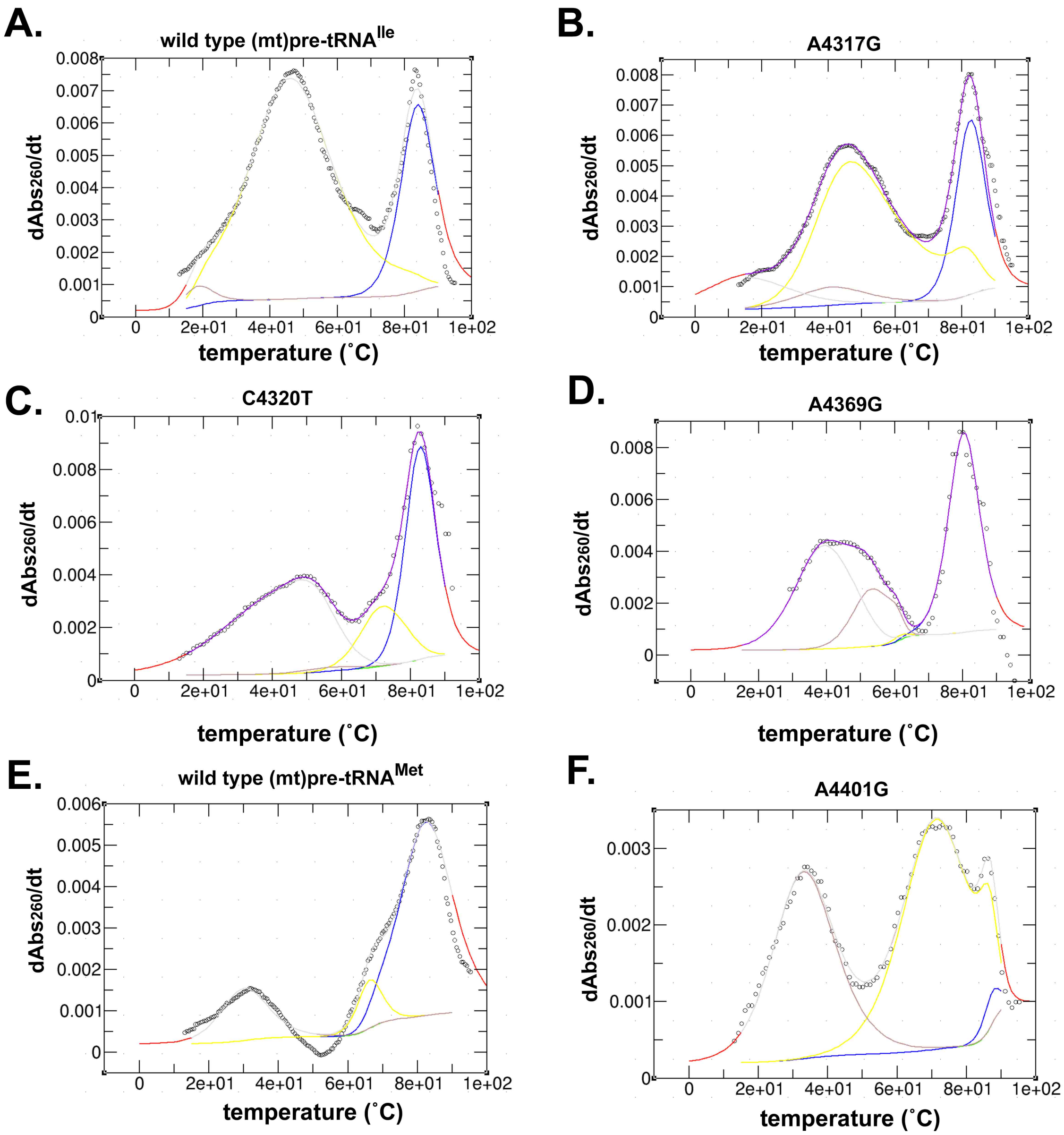


Supplemental Figure 2. Examples of fitting data from UV melting experiments by fitUVData.py and Global Melt Fit [33, 34] for different (mt)pre-tRNAs.
